# Exploring Brain State Changes Through Reaction Time in a Simultaneity Judgment Task: A Pilot Study on Multisensory Integration

**DOI:** 10.1101/2025.09.04.674205

**Authors:** Andrew Jeyathasan, Swati Banerjee

## Abstract

**Problem statement:** Multisensory Integration (MSI), the brain’s ability to merge information from different sensory modalities, relies on the Temporal Binding Window (TBW)—the interval during which stimuli are most likely to be perceived as simultaneous. Beyond its role in perception, MSI may also reflect changes in brain state as task difficulty evolves. In this pilot study, we aimed to explore whether behavioral metrics, particularly reaction time (RT), can capture these brain state changes during a Simultaneity Judgment (SJ) task. Understanding this relationship is critical for developing adaptive Brain-Computer Interfaces (BCIs) that rely on real-time cognitive and perceptual feedback.

**Approach:** To assess how temporal alignment influences both perception and underlying brain state, we employed a Two-Alternative Forced Choice (2AFC) SJ task using audiovisual stimuli. After estimating individual simultaneity thresholds, we implemented an adaptive task that gradually increased difficulty by manipulating the stimulus onset asynchrony (SOA) closer to the TBW. We hypothesized that as perceptual uncertainty increased, RTs would reflect evolving cognitive states, serving as an observable indicator of underlying neural activity shifts.

**Materials and Methods:** Two participants were tested with consistent auditory (500 Hz tone) and visual (faded white circle) stimuli. The TE phase used a staircase procedure to determine personalized TBWs. In the subsequent adaptive test phase, SOAs were progressively modulated around these thresholds to induce varying levels of perceptual ambiguity. Behavioral responses were analyzed using psychometric curve fitting, confusion matrices, and detailed RT evolution across trials.

**Results and Discussion:** Participant 1 showed clear RT modulation near the TBW, suggesting heightened cognitive load and decision uncertainty—a potential signature of brain state transitions. In contrast, Participant 2 exhibited less sensitivity, possibly due to suboptimal thresholding. These preliminary findings suggest that RT holds promise as a behavioral correlate of perceptual state; however, further refinement and standardization of the task are needed before it can be reliably scaled to larger and more controlled studies of MSI-linked brain dynamics.

## I. Introduction

Advances in Brain-Computer Interfaces (BCIs) have opened new avenues for direct communication between the brain and external devices, transforming how humans interact with technology. The success of these systems hinges not only on sophisticated signal acquisition and processing methods, but also on the brain’s remarkable ability to interpret and respond to multisensory cues in real time. Multisensory integration (MSI)—the process by which the brain combines information from different sensory modalities—plays a pivotal role in everyday tasks, from navigating complex environments to mastering skills that require the coordination of visual, auditory, and tactile inputs [1]. This capacity for merging diverse sensory signals is fundamental to immersive technologies, which strive to create seamless and engaging user experiences [2].

Central to MSI is the concept of temporal binding, wherein the brain fuses stimuli that occur close together in time into a unified percept [**?**]. The temporal binding window (TBW) defines the interval during which multisensory signals are most likely to be integrated, and its width can vary depending on stimulus properties, individual differences, and task demands. When sensory inputs are temporally aligned within the TBW, neural responses are synchronized, leading to enhanced perceptual accuracy and faster reaction times. Conversely, misaligned stimuli are processed separately, which can increase cognitive load and reduce system usability.

For BCIs, leveraging temporal binding is crucial for optimizing user performance and experience. Rapid and accurate responses to system cues—often presented through multiple modalities—depend on the brain’s ability to integrate these signals efficiently. Understanding how temporal alignment influences MSI can inform the design of BCI protocols that minimize cognitive overload and maximize real-time control, especially in complex or immersive environments.

In this study, we investigate the impact of temporal alignment on multisensory integration within a BCI context. Using a simultaneity judgment task, we examine whether manipulating the timing of audiovisual stimuli induces measurable changes in brain state, beginning with behavioral metrics such as reaction time. By exploring the boundaries of the temporal binding window and its effects on perception and performance, our work aims to advance the development of more intuitive and effective human-computer interfaces.

## II. Related work

The human brain is an intricate network of billions of neurons communicating through complex pathways. This communication is essential for cognition, movement, sensory perception, and overall neurological function. Past research related to tactile-visual substitution systems has explored the conversion of visual stimuli into tactile sensations, therefore laying the groundwork for sensory augmentation in assistive technologies [3], [4] to overcome the sensory overload in the visual cortex. Studies by Collins [5] have demonstrated the feasibility of inferring visual characteristics from tactile stimuli, underscoring the potential for cross-modal information transfer. This energy conversion phenomenon, where light energy is translated into mechanical or electrical energy, forms the basis of tactile-visual substitution systems [6]. Leveraging this principle, researchers have integrated electro tactile feedback into myoelectric hand prostheses, offering users enhanced sensory perception and control [7]–[9]. Such advancements promise to improve prosthetic devices’ functionality and usability, empowering users with more intuitive and immersive sensory experiences. Oh, Yoon, and Park [10] presented a study to prove that localized electro tactile feedback outperforms visual feedback in a Virtual-reality-based table tennis game. However, this approach remains to be very narrow in terms of applicability. In the following sections, we will explore how different modality can be linked together to create this cross-modal information transfer.

### A. Cross-modal Correspondence

When two or more sensory modalities are presented together, they can influence each other in complex and meaningful ways. Each sense captures different aspects of the environment, yet these modalities can often convey similar or complementary information. This overlap suggests that the effectiveness of MSI depends on how similar or complementary the information is across senses, we then talk about Cross-Modal Correspondences (CMC). Cross Modal correspondence refers to the tendency for normal observers to match distinct features across different sensory modalities [11]. For instance, the association between a sound pitch and size of the object producing the sound [12]. Such correspondences are thought to be supported by shared or interacting brain regions that integrate multisensory input, allowing the brain to form a coherent representation of the environment [13].

CMC across the presented stimuli is essential for effective multisensory integration. Without proper CMC, the brain may struggle to combine inputs from different modalities, which can result in increased reaction times. This effect is illustrated in the pilot study [14] (see Fig.1), where congruent simultaneous cues led to longer reaction times compared to congruent sequential cues. In sequential condition, the visual cue was presented first, followed by the auditory and tactile cues 500 ms later. However, participants often responded before the onset of the auditory and tactile cues, effectively reacting only to the visual cue. Although the sequential condition was technically trimodal, it functioned as a visual unimodal cue in practice. So, using trimodal simultaneous cues leads to an increase in reaction time compared to a visual unimodal cue. This is a clear example of sensory overload, where the brain is unable to process all the information presented simultaneously, leading to higher reaction times and decreased performance.

**Fig. 1:**
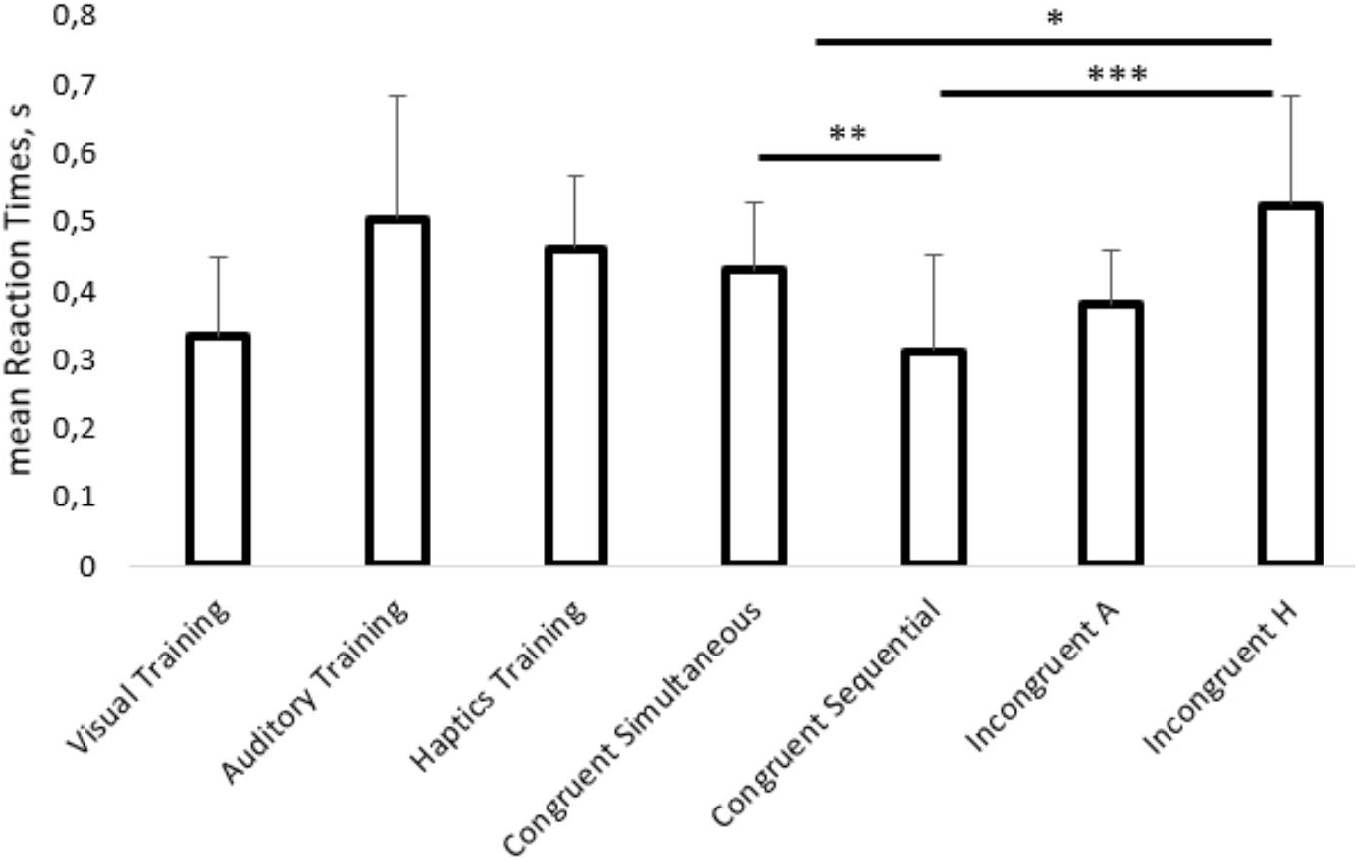
Mean reaction times and accuracies in experiment 1 across different tasks, *p<0.05, **p<0.01, ***p<0.001 in paired t-test. Outliers were excluded from the reaction times. A - auditory, H - haptics. Adapted from Shumkova, Sabir, and Banerjee [14].

Different types of CMC can be used to induce MSI such as location, temporal, semantic, familiarity, etc. Temporal alignment, or the simultaneity of multimodal stimuli is important for effective multisensory integration [15]. For instance, a tactile stimulus on the hand and a visual stimulus on a screen are more likely to be perceived as related when they occur at almost the same time [16], a phenomenon referred to as the Temporal Ventriloquist Effect.

To address this, the concept of the TBW has been introduced—referring to the time interval within which stimuli from different modalities are likely to be integrated as a single perceptual event. Within this window, small temporal discrepancies between modalities can be tolerated, allowing for successful multisensory fusion.

## III. Materials and Methods

### A. Experimental setup

The experiments were conducted in the lab’s office, with the shutters closed to maintain consistent lighting and all doors and windows shut to minimize external distractions. Participants were seated comfortably in front of a computer screen, with their hands resting on a table. The visual stimuli were presented on a 22-inch HP Compaq LA2205wg monitor, while auditory stimuli were delivered through DACOMEX AH760-U headphones. The experimental setup photo is represented in Fig.2.

**Fig. 2:**
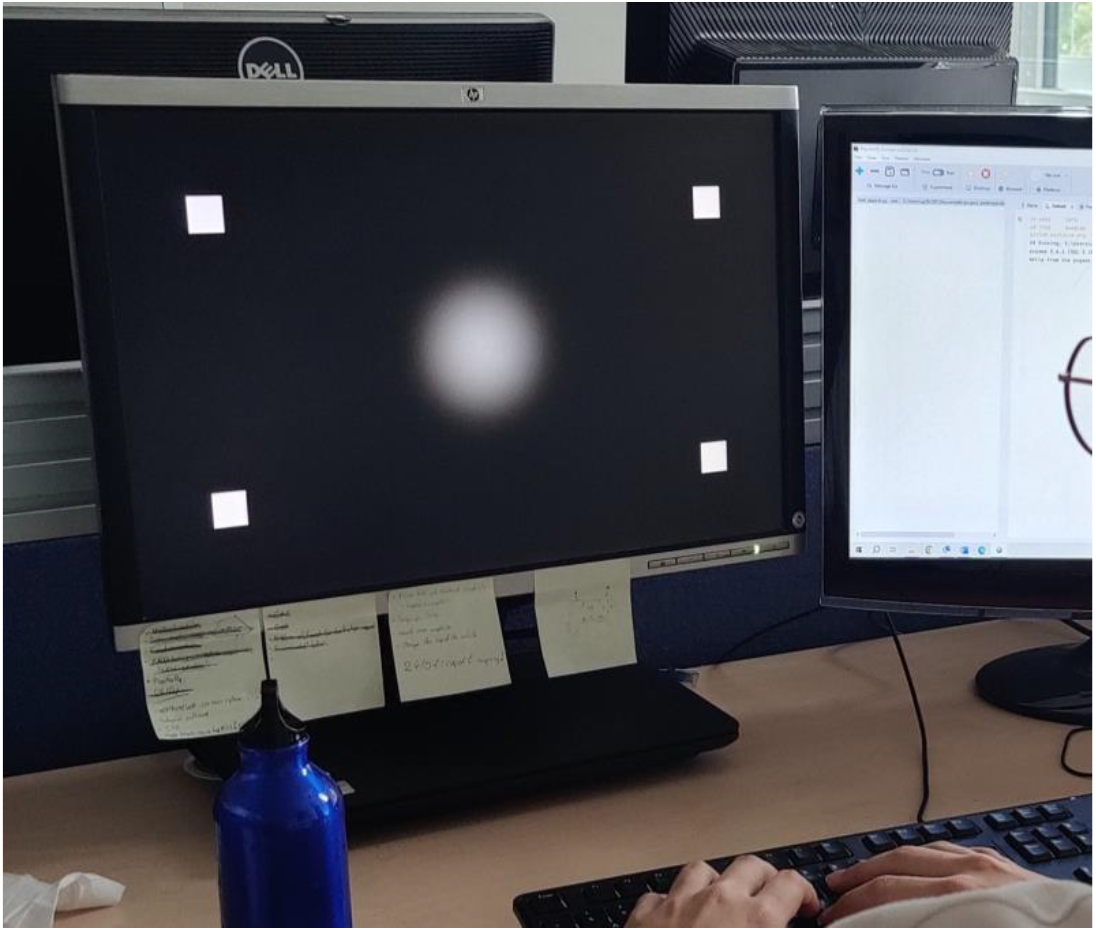
Experimental setup for the multisensory integration tasks.

### B. Assessement and Quantification of MSI

Before delving into the experimental design, it is essential to establish a clear understanding of how to assess and quantify MSI. This section will outline the various methods and metrics used to evaluate MSI, including both behavioral and neural response measures. This understanding will serve as a foundation for the subsequent tasks’ choice and experimental design.

#### 1) Reaction Time

Reaction time (RT) is a widely used behavioral measure in cognitive psychology and neuroscience. It refers to the time taken by an individual to respond to a stimulus, typically measured from the onset of the stimulus to the initiation of a response. RT is often used as an indicator of cognitive processing speed and can provide insights into the efficiency of information processing in the brain. We determined that RT should be a key measure in our experiments, as it provides valuable insights into the efficiency of cognitive processing during multisensory integration. Evidence from a pilot study on the integration of Haptic, auditory and visual (HAV) cues [14] demonstrated that RT was significantly influenced by both the congruency and timing of the presented stimuli (see Fig.1). These findings highlight the importance of considering RT as a sensitive indicator of how different sensory cues are integrated in the brain.

### C. Simultaneity Judgment Task

Perceptual judgment tasks are widely used in cognitive neuroscience to assess how individuals interpret, compare, or discriminate sensory stimuli under varying conditions. These tasks provide a controlled framework for probing the mechanisms by which the brain transforms raw sensory input into meaningful percepts, allowing researchers to isolate specific cognitive processes such as attention, memory, or decision-making [17]. Among these, simultaneity judgment (SJ) tasks are particularly well suited for exploring MSI, as they require participants to evaluate whether stimuli from different sensory modalities occur at the same time. This temporal evaluation directly taps into the brain’s ability to synchronize sensory inputs, a core feature of MSI. By manipulating the timing of cross-modal stimuli and analyzing perceptual responses, SJ tasks enable precise investigation of the TBW [18], [19].

In the following task, we use a Two-alternative forced choice (2AFC) task to investigate the effects of temporal alignment on MSI developed by Binder [20].

#### 1) Stimulation

In the SJ task participants are asked to judge whether two stimuli are presented simultaneously. For this task, we used a faded white circle as the visual stimulus and a 500 Hz tone as the auditory stimulus (see Fig.3). These two stimuli were selected based on the study by Binder [20], although they do not elicit any CMC. However, since our focus is on the effect of temporal alignment on multisensory integration, using two simple stimuli is sufficient to investigate this phenomenon.

**Fig. 3:**
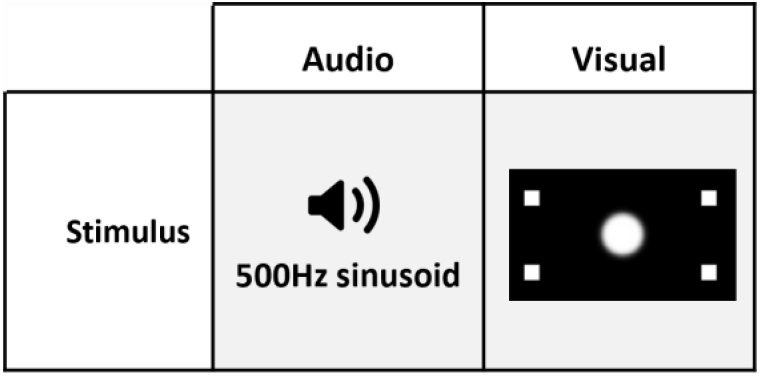
Stimuli used in the SJ tasks.

**Fig. 4:**
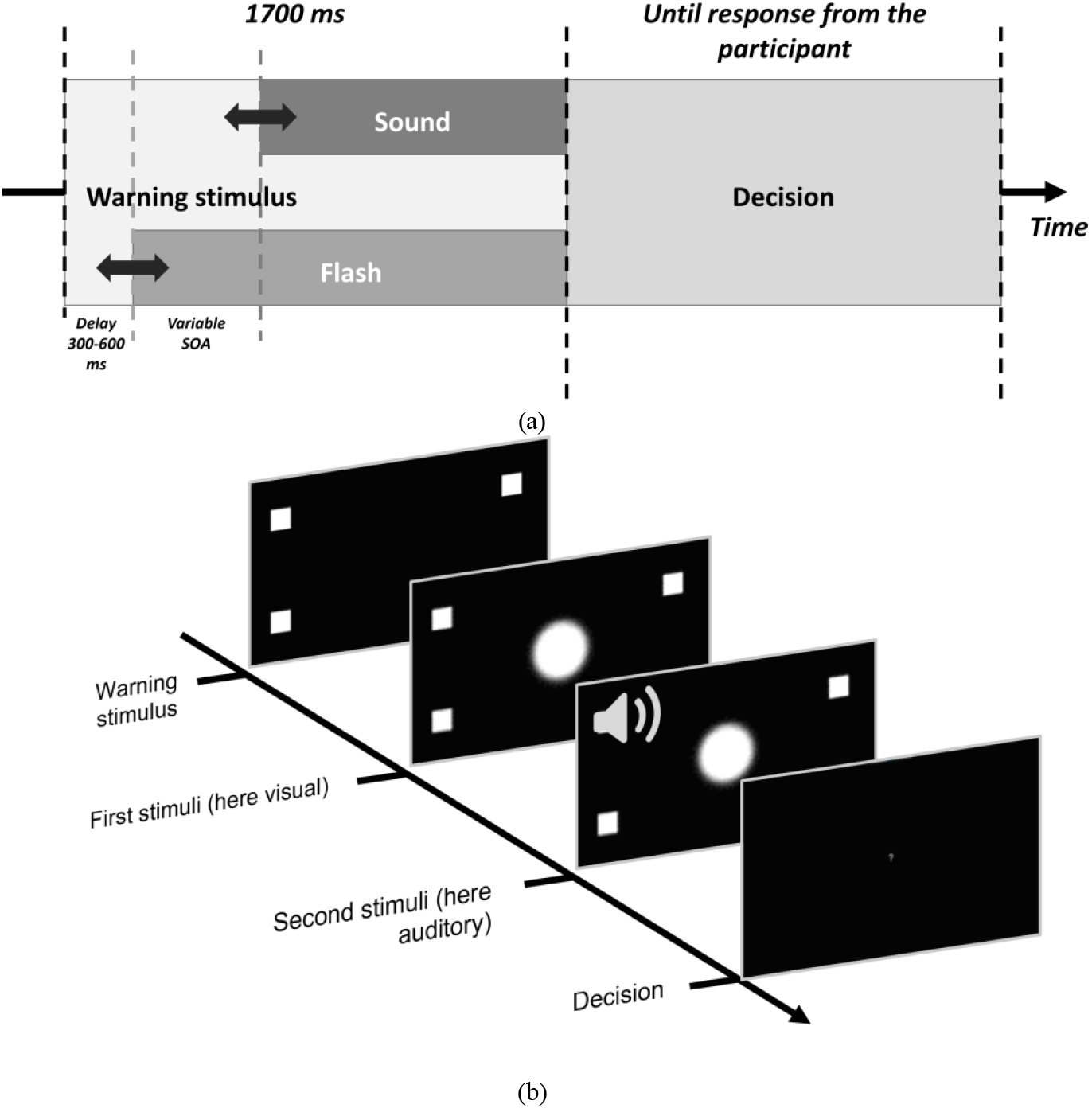
Illustration of the timeline during the SJ task. (a) shows the timing of the stimuli presentation, (b) shows the screen response during the task. Adapted from Binder [20].

#### 2) Experimental paradigm

During the experimental phase, participants performed the SJ task, interleaved with rest periods. At the beginning of each trial, a warning stimulus was displayed—four gray squares surrounding the location of the upcoming visual stimulus. After a randomly selected delay, the auditory and visual stimuli were presented with a specific stimulus onset asynchrony (SOA). SOA values used during the training, test and increase difficulty phase were based on the individual simultaneity thresholds determined during Threshold estimation (TE) phase. There were six possible SOA values:

- Equal to simultaneity threshold
- Half of the threshold value
- Twice of the simultaneity threshold value

Both for sound-first and flash-first trials. Probability of each SOA value was identical and was 0.1667 for each trial (see Fig.5). Presentation of both stimuli terminated simultaneously, and then a message was displayed at the center of the screen to tell his decision (“Did the stimulus happen simultaneously (press “y”) or not (press “n”)”) and reminding which key to press. The message disappeared when subject responded. The task is repeated 6 times and for each repetition, six experimental trials were presented, and the rest duration between the repetitions was 7s.

**Fig. 5:**
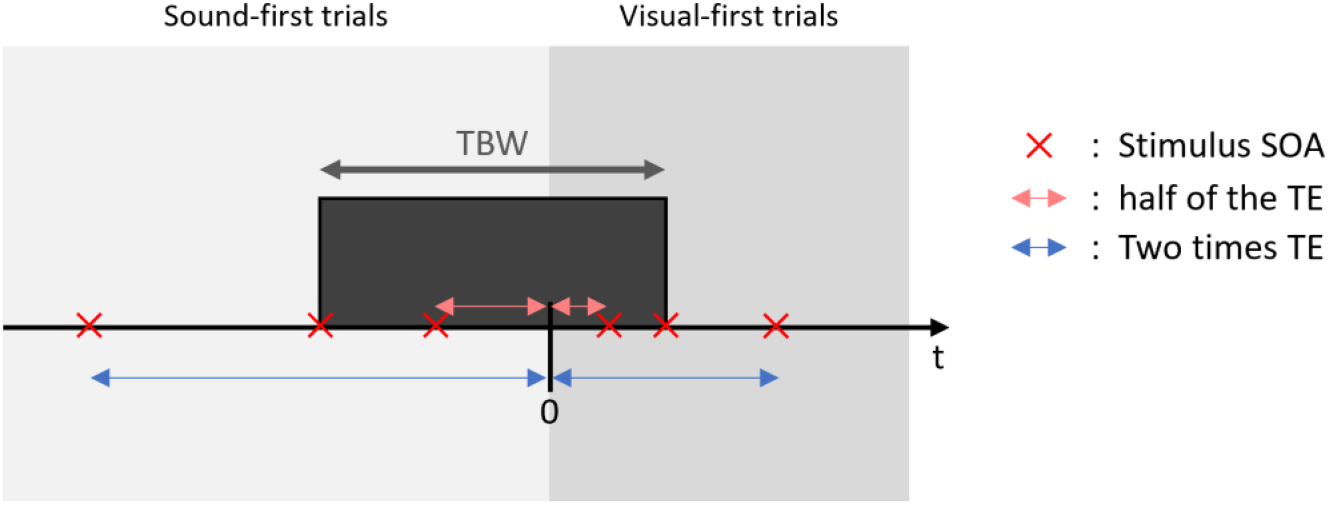
Illustration temporal location of the six SOA values.

#### 3) Thresholding procedure

During the first phase, the TE phase, the individual simultaneity thresholds were determined for each subject. Simultaneity threshold is the point on the psychometric curve where there is an equal probability of the ‘simultaneous’ and ‘non-simultaneous’ response. Thus, the range between simultaneity thresholds for ‘sound-first’ and ‘flash-first’ indicates the SOA values for which the audiovisual pairs are more often judged as synchronous than not (see Fig.5), refered as TBW. An interleaved staircase procedure was used to determine simultaneity thresholds, and it consisted of four staircases with the instruction to judge simultaneity of the presented audiovisual pairs. The SOA values listed below are expressed as the delay of the sound stimulus onset. The initial SOA values of the four staircases were −250 ms, 0 ms for the two staircases with the leading sound stimulus (sound-first trials), and 0 ms, 250 ms for the two staircases with the leading visual stimulus (flash-first trials) (see Fig.6).

**Fig. 6:**
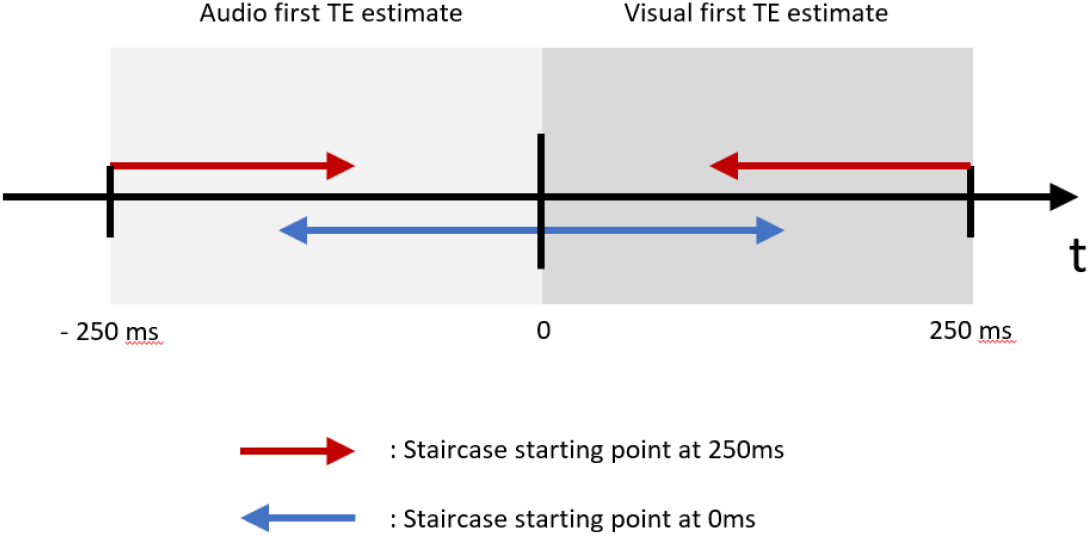
Illustration of the staircase procedure. The initial SOA values of the four staircases were −250 ms, 0 ms for the two staircases with the leading sound stimulus (sound-first trials), and 0 ms, 250 ms for the two staircases with the leading visual stimulus (flash-first trials).

The simultaneity thresholds were calculated separately for sound-first and flash-first trials, as means of SOA values of the last decisions in each of two staircases.

By determining each participant’s individual TBW, this task enables us to compare behavioral and neural responses in conditions where stimuli fall within the TBW—allowing for the ventriloquist effect—and outside the TBW, where such integration is less likely to occur.

#### 4) Adaptive test phase

In this phase, there are 6 SJ tasks with 6 trials each using the same design as the Experimental paradigm but using varying SOA values In fact, to increase the difficulty of the task, we decided to give to the participant, stimuli that are getting closer to the TE (see Fig. 7) so that it is harder for the participant increasing therefore his attention. To make it gradually more difficult, we decided to calculate the new SOA values as follows:

**Fig. 7:**
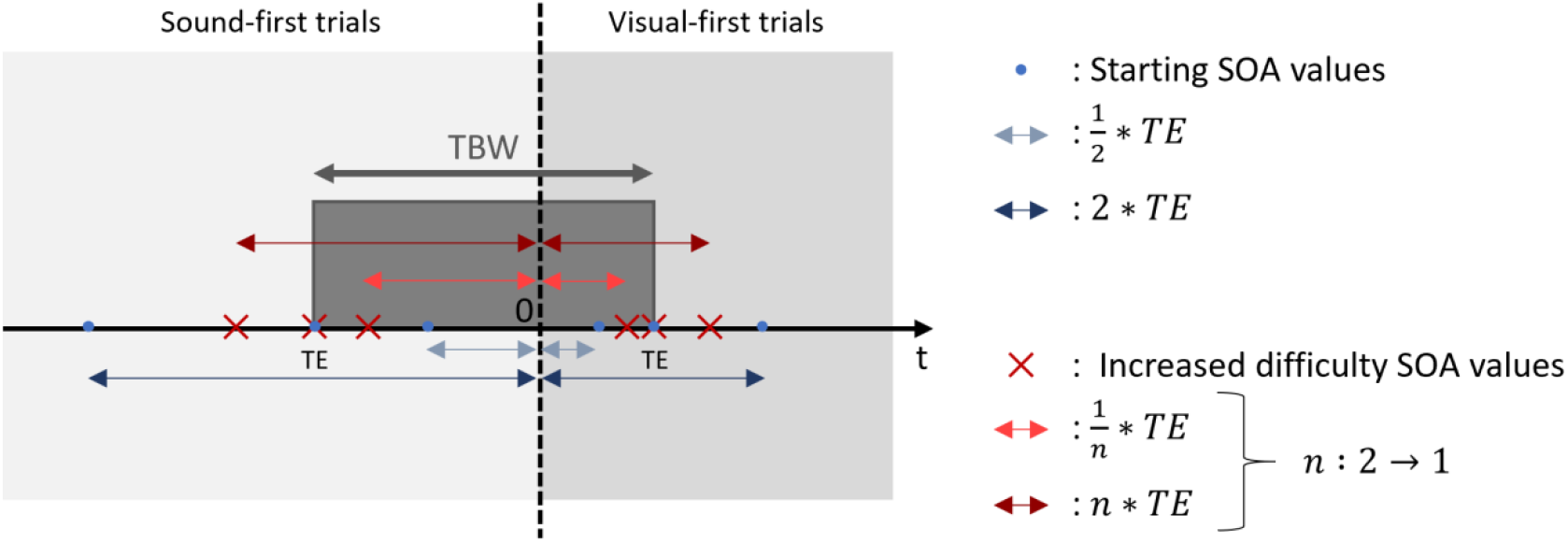
Illustration of the changing SOA during the SJ task adaptive test phase.

- n times the simultaneity threshold : *n* ∗ *TE*
- One over n times the simultaneity threshold : 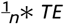

We use this formula both for sound-first and flash-first trials TE, with *n* decreasing from 1.925 to 1.1 in steps of 0.15 every block.

#### 5) Behavioral data analysis

Fitting with a psychometric curve

## IV. Results

### A. Staircase convergence

The staircase procedure for determining individual simultaneity thresholds converged successfully for both participants, as shown in Figures 8a and 8b. The convergence of the staircase procedure is indicated by the stabilization of the SOA values across trials, with the last few trials showing minimal fluctuations. This suggests that both participants were able to reliably judge the simultaneity of the audiovisual stimuli, allowing for accurate estimation of their individual simultaneity thresholds. The final SOA values for Participant 1 were −0.072 seconds for auditory-first and 0.126 seconds for visual-first, while Participant 2’s thresholds were −0.132 seconds for auditory-first and 0.228 seconds for visual-first. These values indicate that both participants have a well-defined TBW, which will be used in subsequent analyses to assess their behavioral and neural responses during the adaptive test phase.

**Fig. 8:**
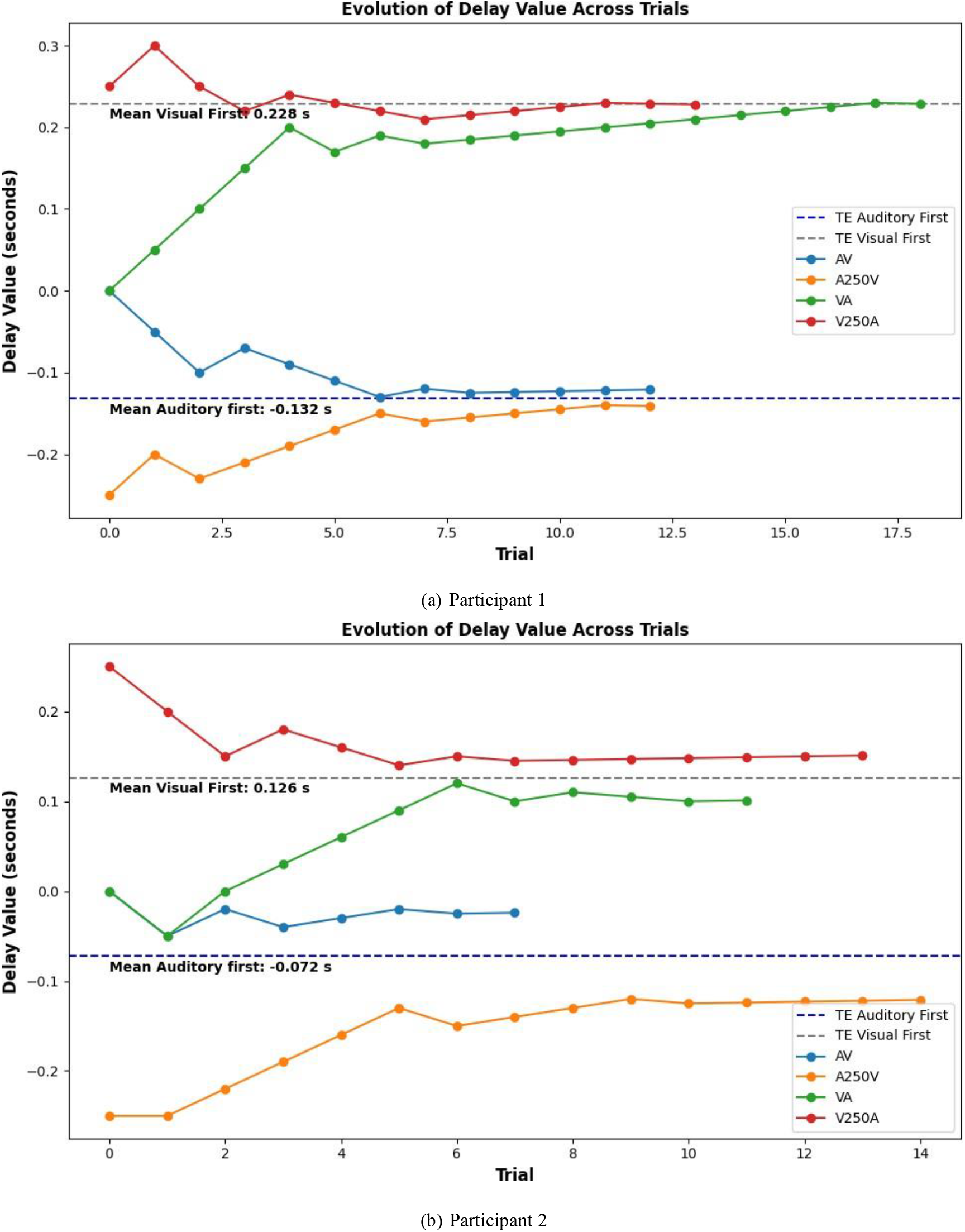
Staircase procedure results for (a) Participant 1 and (b) Participant 2. The x-axis represents the trial number, while the y-axis shows the SOA in seconds. The dashed line indicates the estimated simultaneity threshold.

### B. Psychometric curves

The psychometric curve serves as a crucial verification tool for the proper functioning of the task. By plotting the proportion of “Simultaneous” responses against varying stimulus onset asynchronies (SOA), it allows us to assess whether participants are reliably perceiving and responding to the temporal alignment of audiovisual stimuli. A well-defined curve indicates that the task is effectively capturing the expected perceptual judgments, confirming that the experimental design and stimulus presentation are working as intended. This verification ensures the validity of subsequent analyses and interpretations based on the collected behavioral data.

The psychometric curves shown in Figures 9a and 9b illustrate the proportion of “Simultaneous” responses as a function of SOA for two participants during the thresholding phase. Both curves display a characteristic bell-shaped profile, with the highest probability of simultaneity judgments occurring near zero SOA, and a gradual decline as SOA increases in either direction. This pattern confirms that participants are sensitive to temporal alignment and that the task reliably captures individual simultaneity thresholds.

**Fig. 9:**
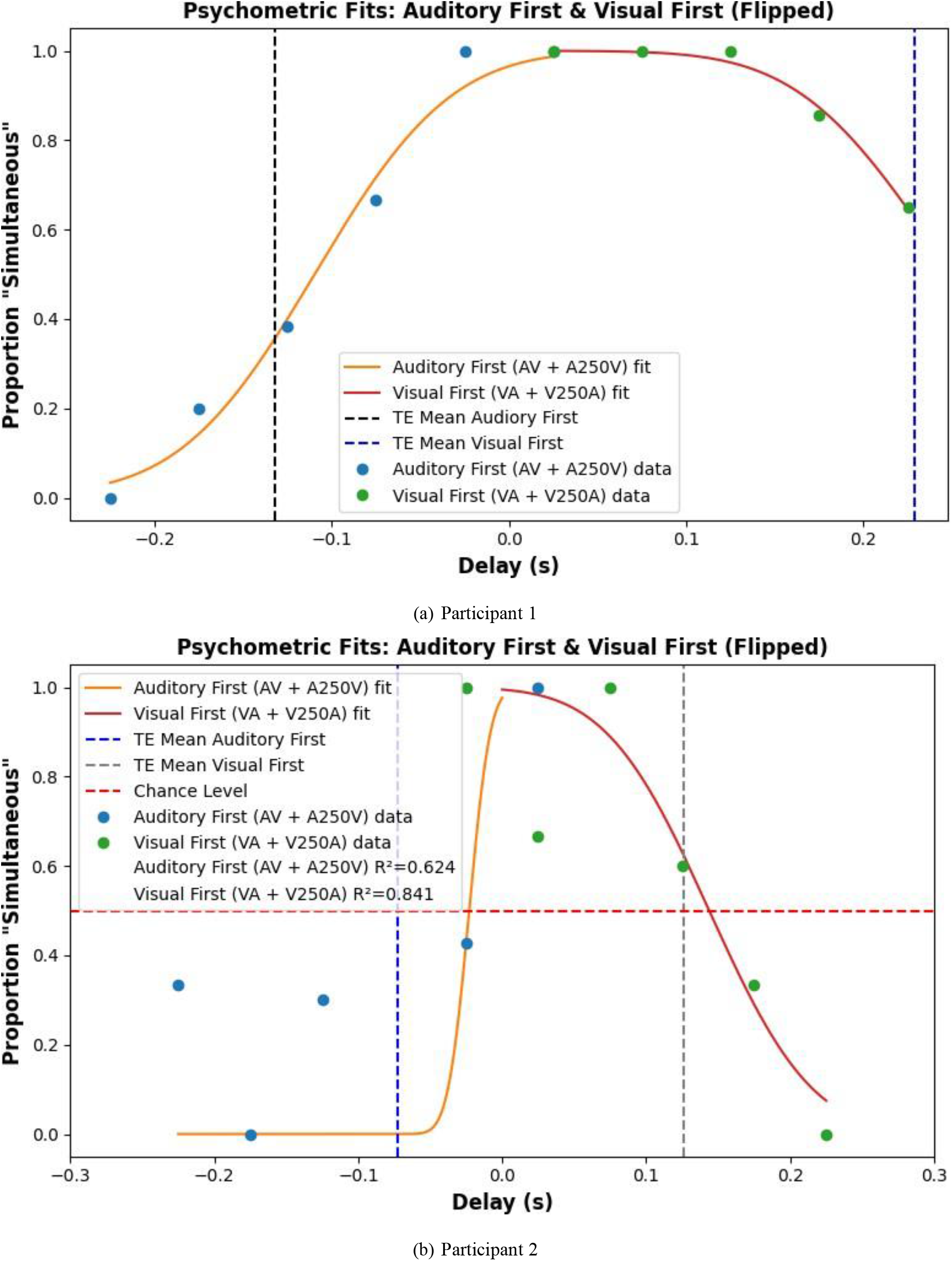
Psychometric curves obtained from the staircase procedure for (a) Participant 1 and (b) Participant 2. The x-axis represents the SOA in seconds, while the y-axis shows the proportion of “Simultaneous” responses.

When comparing the two curves, a notable difference emerges in the quality of the psychometric fit. For Participant 1, the curve closely follows the expected bell-shaped distribution, with a high coefficient of determination (*R*^2^ = 0.984 for auditory-first and *R*^2^ = 0.989 for visual-first), indicating that the model explains most of the variance in the data. In contrast, Participant 2’s curve shows much greater variability in responses and a less precise fit to the model, as reflected by lower *R*^2^ values (*R*^2^ = 0.624 for auditory-first and *R*^2^ = 0.841 for visual-first). This increased variability suggests that the task is not yet sufficiently well defined or standardized, and that the current protocol may not consistently capture the intended perceptual judgments across individuals. Such variability is expected at this stage, as these measurements are part of an initial pilot study aimed at exploring and refining the validity of the task before broader application.

The quality of the psychometric fit is further illustrated in Figures 10a and 10b, which show the psychometric curves obtained from the increasing difficulty phase for two participants. For Participant 1, the psychometric curves remain well defined, with high coefficients of determination (*R*^2^ = 0.883 for auditory-first and *R*^2^ = 0.996 for visual-first), indicating that the model continues to explain most of the variance in the data even as the task becomes more challenging. In contrast, Participant 2’s curves are highly variable and essentially unusable in this phase, as reflected by very low *R*^2^ values (*R*^2^ = 0.030 for auditory-first and *R*^2^ = 0.634 for visual-first). This stark difference highlights how the psychometric curve serves as a valuable tool for assessing the reliability and usability of behavioral data: while Participant 1’s data remain interpretable, the poor fit for Participant 2 suggests that their responses do not consistently reflect the underlying perceptual process, possibly due to increased task difficulty or lack of engagement. Thus, examining the quality of the psychometric fit is crucial for determining which data can be meaningfully analyzed in multisensory integration studies.

**Fig. 10:**
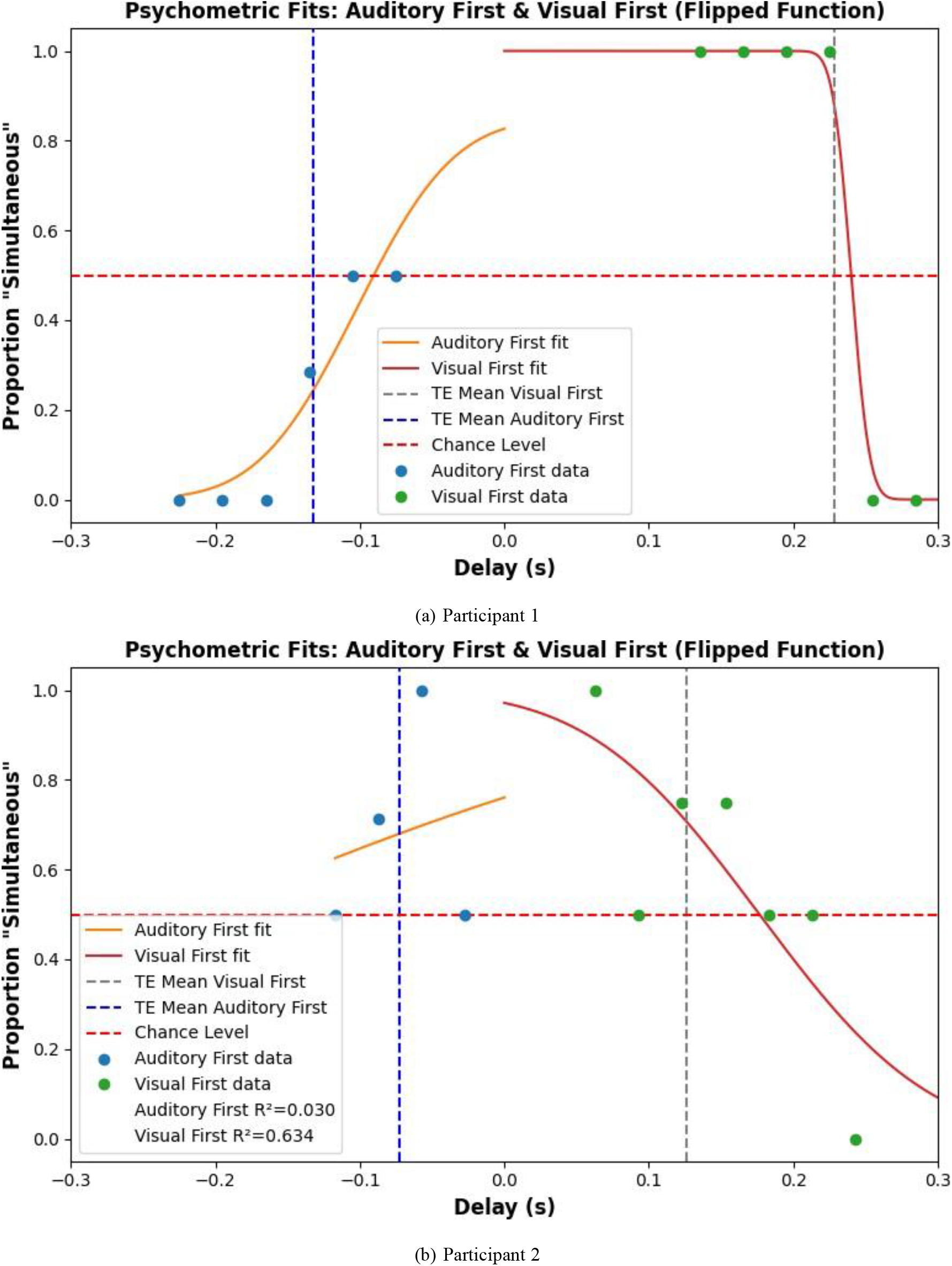
Psychometric curves obtained from the increasing difficulty phase for (a) Participant 1 and (b) Participant 2.

### C. Accuracy

The confusion matrices presented in Figures 11a and 11b provide a clear visualization of the accuracy achieved by each participant during the test phase for the clear conditions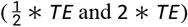. The high concentration of responses along the diagonal indicates that participants were able to correctly identify the simultaneity or non-simultaneity of the presented stimuli with a high degree of reliability. This pattern demonstrates that the task design effectively enables participants to discriminate between different stimulus conditions, supporting the validity of the simultaneity judgment paradigm. The low proportion of off-diagonal responses further suggests minimal confusion between classes, highlighting the robustness of the experimental protocol in eliciting accurate perceptual judgments.

**Fig. 11:**
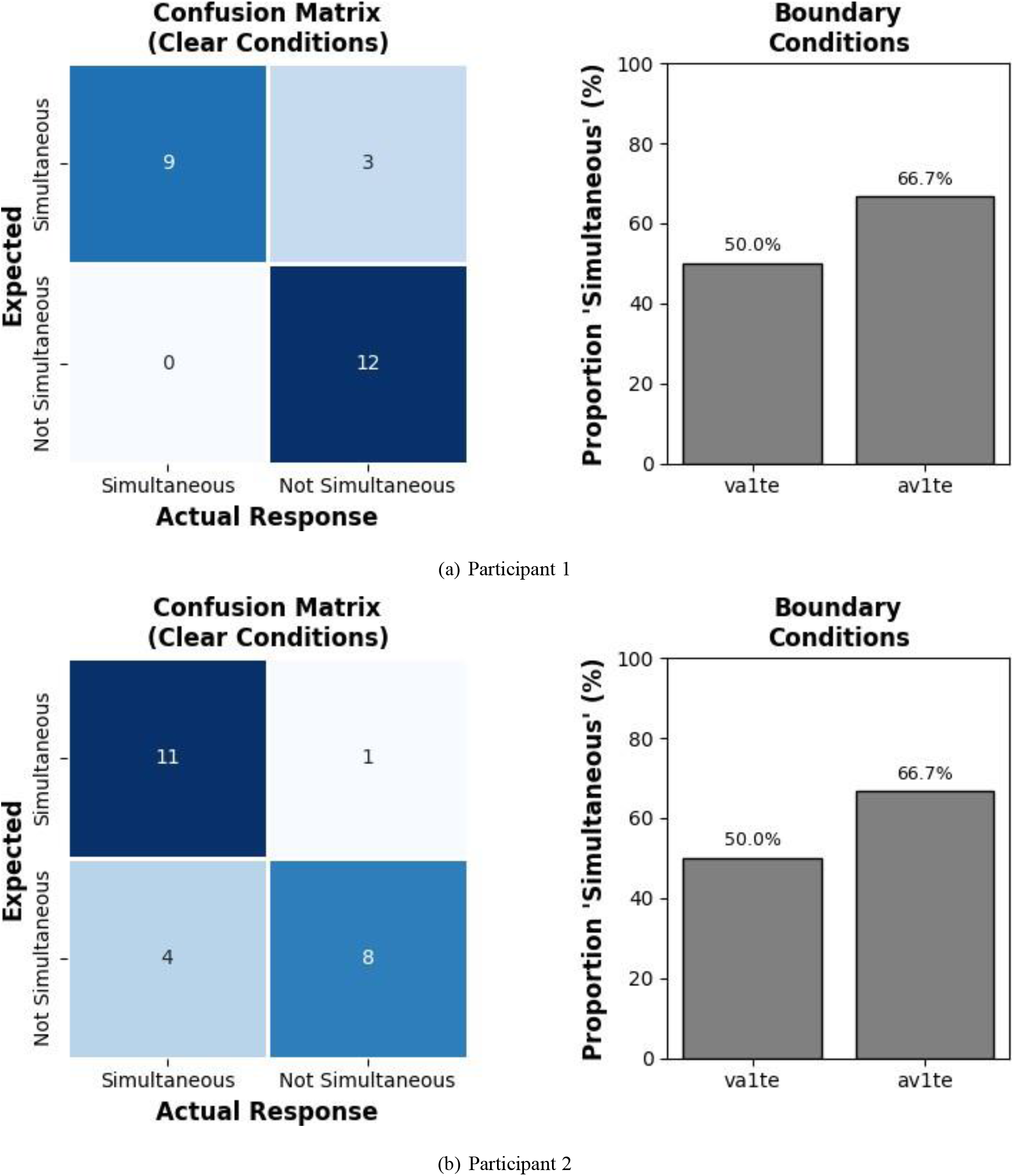
Confusion matrices for the test phase of (a) Participant 1 and (b) Participant 2. The x-axis represents the predicted class, while the y-axis shows the true class. The color intensity indicates the proportion of responses for each combination of true and predicted classes.

For stimuli presented with SOA values at the border of the individual TBW (1 ∗ *TE*), the proportion of “Simultaneous” responses approaches chance level, typically around 50%. This reflects the inherent uncertainty in perceptual judgments when the temporal separation between auditory and visual stimuli is near the threshold for integration. At these SOA values, participants are equally likely to judge the stimuli as simultaneous or non-simultaneous, indicating that the task effectively captures the transition point between integrated and segregated percepts. This pattern further validates the use of the psychometric curve to quantify individual differences in temporal sensitivity and the boundaries of multisensory integration.

The confusion matrices and psychometric curves obtained during the increasing difficulty phase reveal notable differences in participant performance and task validity. For Participant 1, the confusion matrix (Fig. 12a) shows a strong concentration of responses along the diagonal, indicating high accuracy in distinguishing between simultaneous and non-simultaneous stimuli even as the task became more challenging. Correspondingly, the psychometric curve (Fig. 10a) remains well-defined and closely fits the expected bell-shaped profile, with high *R*^2^ values (*R*^2^ = 0.883 for auditory-first and *R*^2^ = 0.996 for visual-first), suggesting consistent and reliable perceptual judgments. In contrast, Participant 2’s confusion matrix (Fig. 12b) displays a more dispersed pattern, with responses less concentrated on the diagonal, reflecting increased confusion and reduced accuracy. This is further supported by the psychometric curve (Fig. 10b), which exhibits high variability and poor model fit (*R*^2^ = 0.030 for auditory-first and *R*^2^ = 0.634 for visual-first), indicating inconsistent responses as task difficulty increased.

**Fig. 12:**
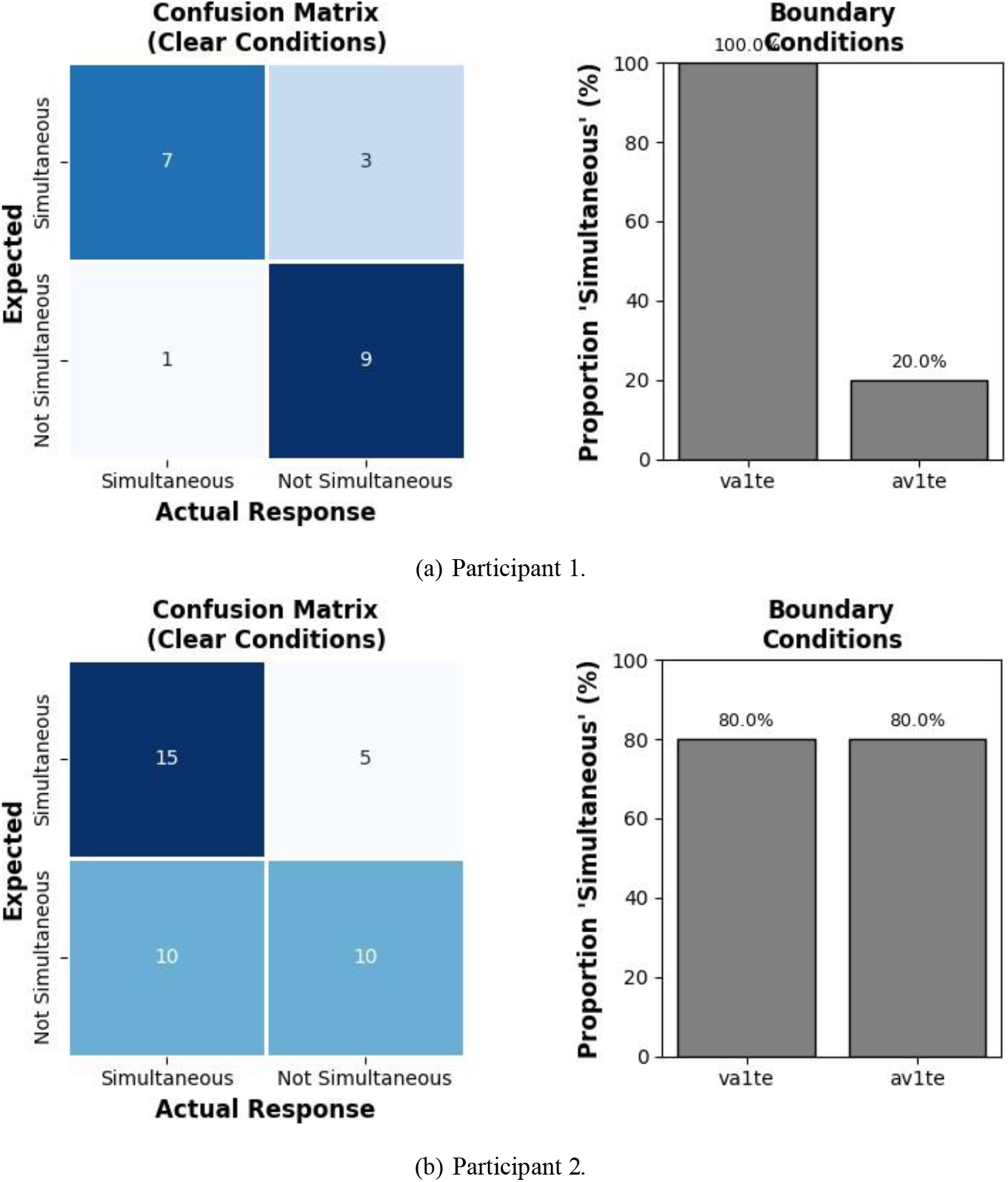
Confusion matrices for the increasing difficulty phase of (a) Participant 1 and (b) Participant 2. The x-axis represents the predicted class, while the y-axis shows the true class. The color intensity indicates the proportion of responses for each combination of true and predicted classes.

These observations suggest that while the task is capable of eliciting precise and interpretable behavioral data for some participants, it may not be sufficiently robust or standardized for all individuals. The strong performance and clear psychometric fit for Participant 1 support the validity of the task under increased difficulty, demonstrating its potential to capture meaningful differences in temporal sensitivity. However, the variability and poor fit observed for Participant 2 highlight the need for further refinement and standardization of the protocol to ensure consistent and reliable measurement across participants. Importantly, the poor accuracy and high variability for Participant 2 may also be explained by a suboptimal threshold estimate for the Temporal Binding Window (TBW). If the thresholding procedure did not accurately capture the true simultaneity threshold, the SOA values used in the adaptive phase may have been inappropriate, resulting in increased uncertainty and inconsistent responses. This underscores the importance of robust and individualized thresholding to ensure valid assessment of multisensory integration and highlights the need for iterative task development in future studies.

### D. Reaction Time

The reaction time (RT) figures reveal clear differences between participants in how RT varies across delay conditions and task difficulty. For Participant 2, RTs remain relatively stable and low across all SOA conditions, both in the binned (see Fig.13) and grouped bar plots (see Fig.14), with only minor fluctuations as the delay changes. This suggests that Participant 2 was able to respond quickly and consistently, regardless of the temporal alignment of the stimuli—even as the task became more challenging. In contrast, Participant 1 shows much greater variability in RT, with pronounced increases in mean RT for certain SOA values, especially as the delay approaches the individual threshold estimate (TE). The bar plots for Participant 1 also display higher mean RTs and larger error bars, indicating both slower and more inconsistent responses across conditions. These observations suggest that while the task elicits robust and reliable RT measurements for some participants, others may experience increased cognitive load or uncertainty as the temporal alignment becomes more ambiguous. For Participant 2, the stable and low RTs across all SOA conditions suggests that the task may not be functioning as intended for this participant. Ideally, reaction times should increase for SOA values closer to the individual threshold (TE), reflecting increased decision difficulty. The absence of this pattern for Participant 2 indicates that the task may not have been sufficiently challenging or that the participant was not sensitive to the temporal manipulation due to a wrong TE. In contrast, Participant 1 shows pronounced increases in mean RT for SOA values near the TE, as expected when the task becomes more difficult and decisions are less clear. This pattern supports the validity of the task for Participant 1 and highlights the importance of observing increased RTs under more ambiguous conditions. Therefore, the lack of RT variability for Participant 1 suggests a limitation in the task’s sensitivity or calibration, emphasizing the need for further refinement to ensure that the task reliably captures temporal discrimination across all participants.

**Fig. 13:**
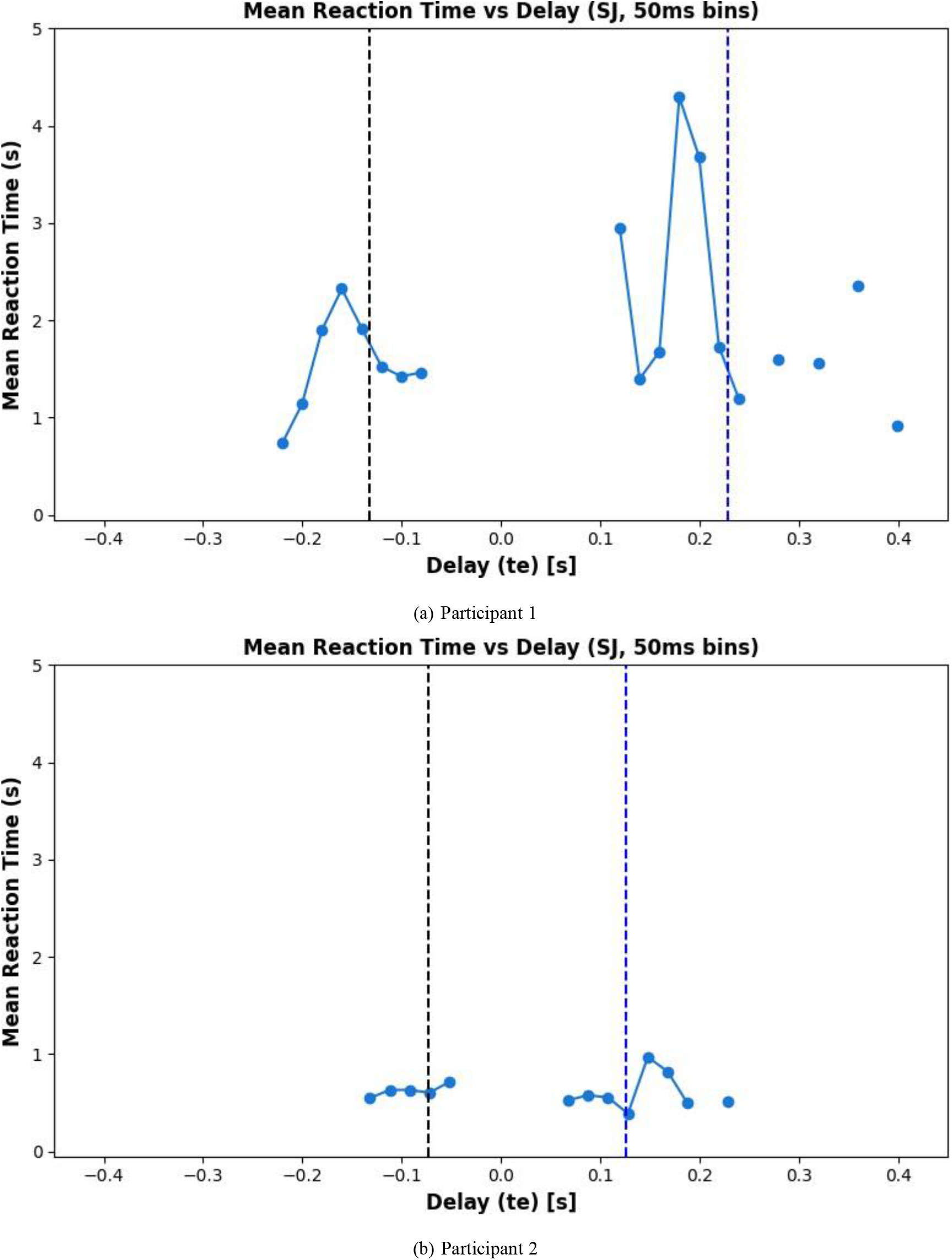
Mean Reaction time in a 20 ms bin during the increasing difficulty phase for (a) Participant 1 and (b) Participant 2. The x-axis represents the SOA in seconds, while the y-axis shows the reaction time in seconds.

**Fig. 14:**
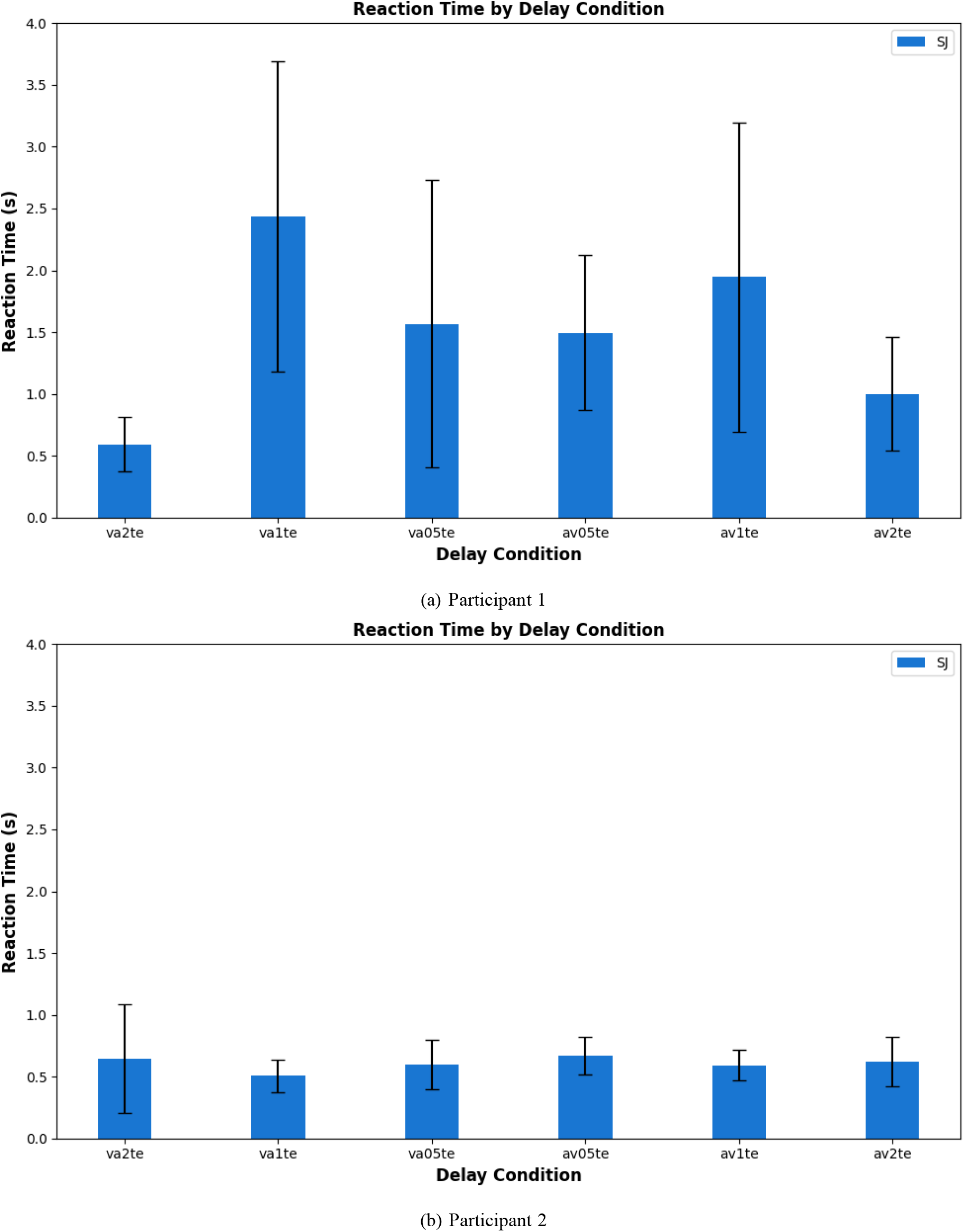
Reaction time during the test phase for each condition and for (a) Participant 1 and (b) Participant 2.

## V. Discussion and conclusion

Talk about the brain state changes and the bayesian model

One area for improvement in the current protocol is the standardization of instructions and procedures across participants. In this study, Participant 1 was not informed that reaction time would be measured, which led them to spend more time analyzing each stimulus and resulted in a more accurate psychometric curve. In contrast, Participant 2 was aware that reaction time was being recorded, prompting faster responses but yielding less precise data for accuracy and psychometric analysis. These inconsistencies highlight the need for a more standardized approach to task instructions and experimental procedures. By ensuring that all participants receive the same information and guidance, future studies can achieve more reliable and consistent results, improving the validity and comparability of multisensory integration findings.

While this study focused on the temporal binding window between visual and auditory stimuli, an important future direction is to extend the protocol to include haptic (tactile) stimulation. By investigating visual-tactile and auditory-tactile combinations, we can gain a more comprehensive understanding of multisensory integration mechanisms. Incorporating haptic cues would allow us to examine how temporal alignment across three modalities—visual, auditory, and tactile—affects perception and reaction time. A promising approach is to adapt the simultaneity judgment task to use trimodal cues, enabling the assessment of temporal binding windows in more complex multisensory environments.

Another important consideration for future studies is the incorporation of noise stimuli, which are prevalent in real-world multisensory environments. By introducing controlled noise across modalities, we can better mimic natural conditions and investigate how the presence of irrelevant or distracting sensory information influences multisensory integration. This approach may enhance the ecological validity of the protocol and provide deeper insights into how the brain maintains robust integration and perceptual accuracy in everyday settings. Together, these extensions would offer valuable perspectives on cross-modal interactions and inform the design of more immersive and effective human-computer interfaces.

## List of Abbreviations

2AFC: Two-Alternative Forced Choice
CMC: Cross-Modal Correspondence
EEG: Electroencephalogram
HAV: Haptic-Audio-Visual
MSI: Multisensory Integration
RT: Reaction Time
SJ: Simultaneity Judgment
SOA: Stimulus Onset Asynchrony
TBW: Temporal Binding Window
TE: Threshold Estimation

## Acknowledgment

The first author is thankful for the funding provided by CEA LIST under the Frugal Brain-HAVSENSE [CEA-SI-2022-004] project initiative.

## References

[1] M. Marucci, G. Di Flumeri, G. Borghini, N. Sciaraffa, M. Scandola, E. F. Pavone, F. Babiloni, V. Betti, and P. Aricò, “The impact of multisensory integration and perceptual load in virtual reality settings on performance, workload and presence,” Scientific Reports, vol. 11, no. 1, p. 4831, Mar. 2021. [Online]. Available: https://www.nature.com/articles/s41598-021-84196-8

[2] N. Curran, “Factors of Immersion,” in The Wiley Handbook of Human Computer Interaction, 1st ed., K. L. Norman and J. Kirakowski, Eds. Wiley, Feb. 2018, pp. 239–254. [Online]. Available: https://onlinelibrary.wiley.com/doi/10.1002/9781118976005.ch13

[3] P. Bach-y Rita, “Late postacute neurologic rehabilitation: neuroscience, engineering, and clinical programs,” Archives of Physical Medicine and Rehabilitation, vol. 84, no. 8, pp. 1100–1108, Aug. 2003.

[4] C. Phunruangsakao, D. Achanccaray, S. Bhattacharyya, S.-I. Izumi, and M. Hayashibe, “Effects of visual-electrotactile stimulation feedback on brain functional connectivity during motor imagery practice,” Scientific Reports, vol. 13, p. 17752, Oct. 2023. [Online]. Available: https://www.ncbi.nlm.nih.gov/pmc/articles/PMC10584917/

[5] C. C. Collins, “Tactile Television - Mechanical and Electrical Image Projection,” IEEE Transactions on Man-Machine Systems, vol. 11, no. 1, pp. 65–71, Mar. 1970, conference Name: IEEE Transactions on Man-Machine Systems. [Online]. Available: https://ieeexplore.ieee.org/document/4081932

[6] M. Isaković, J. Malešević, T. Keller, M. Kostić, and M. Štrbac, “Optimization of Semiautomated Calibration Algorithm of Multichannel Electrotactile Feedback for Myoelectric Hand Prosthesis,” Applied Bionics and Biomechanics, vol. 2019, p. 9298758, Mar. 2019. [Online]. Available: https://www.ncbi.nlm.nih.gov/pmc/articles/PMC6437744/

[7] J. Dong, W. Jensen, B. Geng, E. N. Kamavuako, and S. Dosen, “Online Closed-Loop Control Using Tactile Feedback Delivered Through Surface and Subdermal Electrotactile Stimulation,” Frontiers in Neuroscience, vol. 15, p. 580385, 2021.

[8] G. Chai, H. Wang, G. Li, X. Sheng, and X. Zhu, “Electrotactile Feedback Improves Grip Force Control and Enables Object Stiffness Recognition While Using a Myoelectric Hand,” IEEE transactions on neural systems and rehabilitation engineering: a publication of the IEEE Engineering in Medicine and Biology Society, vol. 30, pp. 1310–1320, 2022.

[9] Y. Han, Y. Lu, Y. Zuo, H. Song, C.-H. Chou, X. Wang, X. Li, L. Li, C. M. Niu, and W. Hou, “Substitutive proprioception feedback of a prosthetic wrist by electrotactile stimulation,” Frontiers in Neuroscience, vol. 17, p. 1135687, 2023.

[10] S. Oh, B.-J. Yoon, and H. Park, “Location-based electrotactile feedback localizes hitting point in virtual-reality table tennis game,” Biomedical Engineering Letters, vol. 14, no. 3, pp. 593–604, May 2024.

[11] L. Schmitz, G. Knoblich, O. Deroy, and C. Vesper, “Crossmodal correspondences as common ground for joint action,” Acta Psychologica, vol. 212, p. 103222, Jan. 2021. [Online]. Available: https://linkinghub.elsevier.com/retrieve/pii/S0001691820305461

[12] M. L. Lowe and K. L. Haws, “Sounds Big: The Effects of Acoustic Pitch on Product Perceptions,” Journal of Marketing Research, vol. 54, no. 2, pp. 331–346, Apr. 2017, publisher: SAGE Publications Inc. [Online]. Available: 10.1509/jmr.14.0300

[13] B. E. Stein, T. R. Stanford, and B. A. Rowland, “Multisensory Integration and the Society for Neuroscience: Then and Now,” The Journal of Neuroscience, vol. 40, no. 1, pp. 3–11, Jan. 2020. [Online]. Available: https://www.ncbi.nlm.nih.gov/pmc/articles/PMC6939490/

[14] D. Shumkova, J. Sabir, and S. Banerjee, “CrossModal Correspondence based MultisensoryIntegration: A pilot study showing how HAV cues can modulate the reaction time,” Submited, 2024.

[15] R. L. Miller, S. R. Pluta, B. E. Stein, and B. A. Rowland, “Relative Unisensory Strength and Timing Predict Their Multisensory Product,” The Journal of Neuroscience, vol. 35, no. 13, pp. 5213–5220, Apr. 2015. [Online]. Available: https://www.jneurosci.org/lookup/doi/10.1523/JNEUROSCI.4771-14.2015

[16] R. A. Stevenson, J. K. Fister, Z. P. Barnett, A. R. Nidiffer, and M. T. Wallace, “Interactions between the spatial and temporal stimulus factors that influence multisensory integration in human performance,” Experimental brain research. Experimentelle Hirnforschung. Experimentation cerebrale, vol. 219, no. 1, pp. 121–137, May 2012. [Online]. Available: https://www.ncbi.nlm.nih.gov/pmc/articles/PMC3526341/

[17] J. V. Baranski and W. M. Petrusic, “The calibration and resolution of confidence in perceptual judgments,” Perception & Psychophysics, vol. 55, no. 4, pp. 412–428, Jul. 1994. [Online]. Available: 10.3758/BF03205299

[18] R. A. Stevenson, S. H. Baum, J. Krueger, P. A. Newhouse, and M. T. Wallace, “Links between temporal acuity and multisensory integration across lifespan,” Journal of experimental psychology. Human perception and performance, vol. 44, no. 1, pp. 106–116, Jan. 2018. [Online]. Available: https://www.ncbi.nlm.nih.gov/pmc/articles/PMC5659980/

[19] V. Van Wassenhove, K. W. Grant, and D. Poeppel, “Temporal window of integration in auditory-visual speech perception,” Neuropsychologia, vol. 45, no. 3, pp. 598–607, Jan. 2007. [Online]. Available: https://linkinghub.elsevier.com/retrieve/pii/S002839320600011X

[20] M. Binder, “Neural correlates of audiovisual temporal processing – Comparison of temporal order and simultaneity judgments,” Neuroscience, vol. 300, pp. 432–447, Aug. 2015. [Online]. Available: https://www.sciencedirect.com/science/article/pii/S0306452215004431

